# Chromosome-level genome assembly of the lemon sole *Microstomus kitt* (Pleuronectiformes: Pleuronectidae)

**DOI:** 10.1101/2025.04.29.651060

**Authors:** Marcel Nebenführ, David Prochotta, Maria A. Nilsson, Menno J. de Jong, Tunca D. Yazici, Fabienne Langefeld, Malambo Muloongo, Helena Woköck, Jakob Jilg, Sina C. Bender, Marvin M. Zangl, Juan M. Ortega Guatame, Kimberley Williams, Moritz Sonnewald, Axel Janke

**Affiliations:** Senckenberg Biodiversity and Climate Research Centre (BiK-F), Frankfurt am Main, Germany; Institute for Ecology, Evolution, and Diversity, Goethe University, Frankfurt am Main, Germany; LOEWE-Centre for Translational Biodiversity Genomics (TBG), Frankfurt am Main, Germany; Centre for Wildlife Genetics, Senckenberg Research Institute and Natural History Museum, 63571, Gelnhausen, Germany; Senckenberg Research Institute, Department of Marine Zoology, Section Ichthyology, Frankfurt am Main, Germany; Department of Aquaculture & Fisheries Management, University of Ibadan, Nigeria

**Keywords:** Lemon sole, Pleuronectidae, HiFi long-read sequencing, Proximity ligation sequencing, genome assembly, genome annotation

## Abstract

**Background:** The lemon sole (*Microstomus kitt*) is a culinary fish from the family of righteye flounders (Pleuronectidae) inhabiting sandy and shallow offshore grounds of the North Sea, the western Baltic Sea, the English Channel, the shallow waters of Great Britain and Ireland as well as the Bay of Biscay and the coastal waters of Norway.

**Findings:** Here, we present the chromosome-level genome assembly of the lemon sole. We applied PacBio HiFi sequencing on the PacBio Revio system to generate a highly complete and contiguous reference genome. The resulting assembly has a contig N50 of 17.2 Mbp and a scaffold N50 of 27.2 Mbp. The total assembly length is 628 Mbp, of which 616 Mbp were scaffolded into 24 chromosome-length scaffolds. The identification of 99.7% complete BUSCO genes indicates a high assembly completeness.

**Conclusions:** The chromosome-level genome assembly of the lemon sole provides a high-quality reference genome for future population genomic analyses of a commercially valuable edible fish.

## Data description

### Background information

The lemon sole (*Microstomus kitt*) is a commercially relevant, demersal and bottom-dwelling flatfish species. It can be found on sandy and gravel substrates of the continental shelf, and it feeds primarily on polychaetes. It can reach lengths of 66 cm but is rarely larger than 40 cm and lives for up to 17 years [1]. Females tend to outnumber the males in a population, while also exhibiting a larger size compared to the male specimens [2].

With an estimated catch of about 11,000 tons per year with an estimated value of 36 million €, it constitutes only a small percentage in the European fishery industry [3]. The lemon sole is exclusively available from wild catches by trawling as no farming methods have been developed yet. Due to strict EU regulations, the population is stable (IUCN: least concern) but has not yet been studied for its genomic variability or population genomics [4].

The lemon sole belongs to the Pleuronectidae (righteye flounders), a family of flatfishes in which both eyes wandering to the right body side during development that also includes the halibut (*Hippoglossus hippoglossus)*. This characteristic makes the lemon sole an interesting species for developmental biology [5,6].

Here, we present the chromosome-level genome assembly of the lemon sole *Microstomus kitt*, which may provide important baseline data for future population genetics analyses, for example as part of monitoring efforts in the fishery industry. The genome assembly was performed throughout a six-week master’s course at Frankfurt Goethe University, Frankfurt am Main, Germany. For a detailed description and outline of the course, see [7].

### Sampling, DNA extraction, and sequencing

We obtained a male specimen of the lemon sole (NCBI: txid106175) during a yearly monitoring expedition to the Dogger Bank in the North Sea (54° 48.271′ N 1° 25.077′ E) in July 2023, with the permission of the Maritime Policy Unit of the UK Foreign and Commonwealth Office. The study complied with the ‘Nagoya Protocol on Access to Genetic Resources and the Fair and Equitable Sharing of Benefits Arising from Their Utilization’. The sample was initially frozen at -20 °C and later stored at -80 °C. We extracted high molecular weight DNA from muscle tissue using the QIAGEN DNeasy Blood & Tissue Kit. The quality and quantity of the DNA were determined using the Qubit Fluorometric Quantification by ThermoScientific and the Agilent 2200 TapeStation system (Agilent Technologies).

For long-read libraries, we followed the protocol for the SMRTbell Express Prep kit v3.0 (Pacific Biosciences - PacBio, Menlo Park, CA, USA) and sequenced it at Bioscientia (Ingelheim, Germany) on the PacBio Revio platform. We acquired a total of 5.6 million reads, yielding 64 Gbp sequencing data, with a mean read length of 11.5 kbp. For scaffolding, we generated proximity ligated DNA from heart tissue using the Arima High Coverage HiC Kit v1 (Arima Genomics, Carlsbad, CA, USA), according to the Animal Tissue User Guide. Subsequently, HiC library preparation was conducted following the protocol of the Swift Biosciences® Accel-NGS® 2S Plus DNA Library Kit. Afterwards, the DNA concentration was measured using the Qubit Fluorometer (Thermo Fisher Scientific, Waltham, MA, USA), and the quality was assessed through fragment size distribution using the TapeStation 4150 (Agilent Technologies). The library was then sequenced on the NovaSeq 6000 platform at Novogene (Cambridge, United Kingdom) using a 150 paired-end sequencing strategy, resulting in an output of 269 million reads, resembling 40 Gbp sequencing data in total.

### Genome size estimation

The genome size estimation of the lemon sole was done by analyzing k-mer frequencies. Therefore, HiFi reads were used to determine the k-mer frequency for K=21, running the software jellyfish v2.3.0 [8] and GenomeScope v2.0 [9]. The genome size was estimated to be around 542 Mbp with a heterozygosity of 1.17 % (Figure 1A).

**Figure 1.**
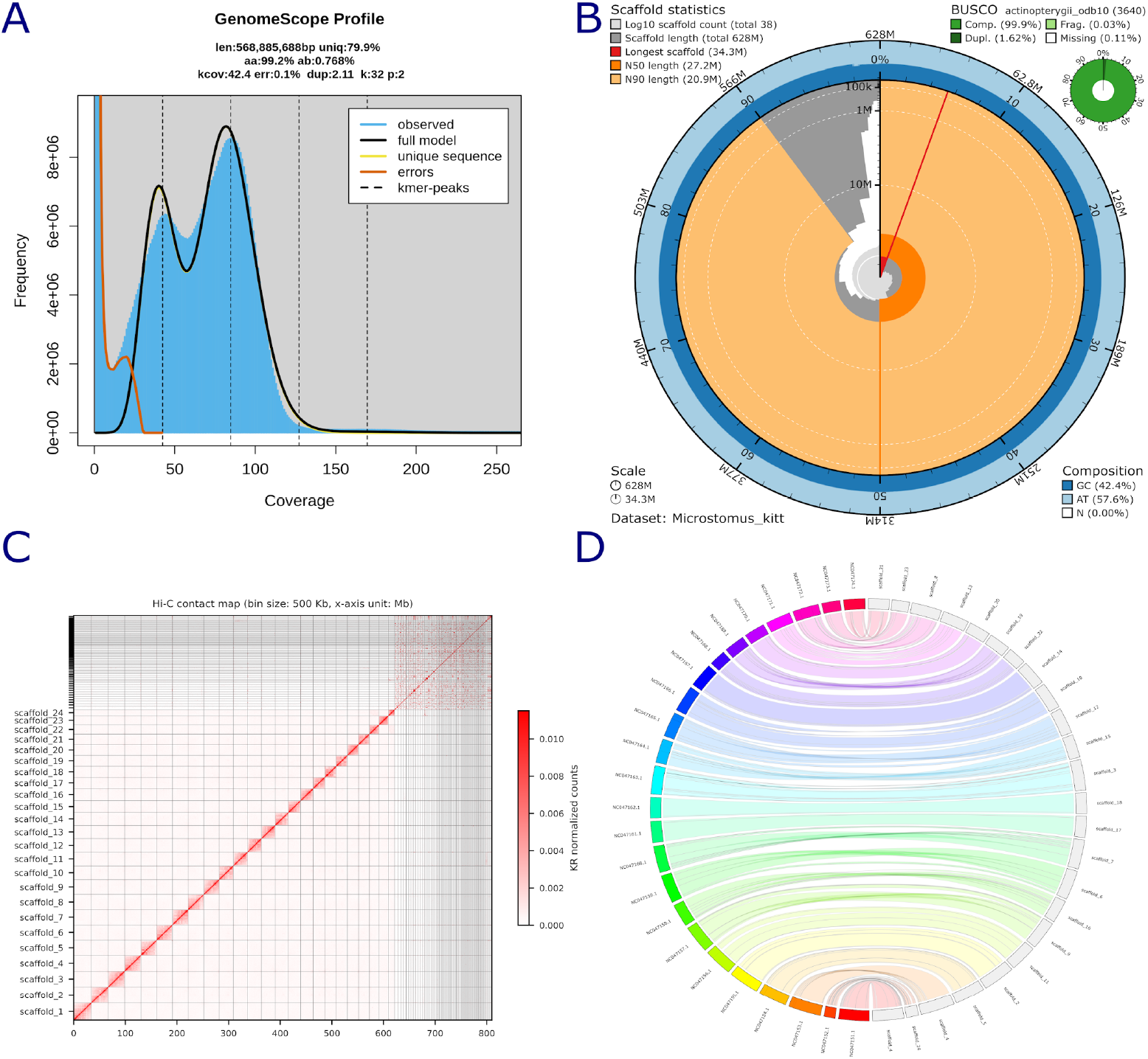
(A) Estimated k-mer distribution calculated with Jellyfish and visualized with GenomeScope as linear plot. (B) Snail plot of the *de novo* genome assembly of the lemon sole showing assembly quality measures. Scaffold statistics are shown by the innermost circle for longest scaffold, N50 and N90 (colors from red to orange). The outer circle (blue) shows GC content. Gene completeness is shown by the green circle. **(C) HiC contact map of the 24 chromosome-length scaffolds and unplaced scaffolds. (D) Whole genome synteny between the final HiC scaffolded chromosome-level assembly of the lemon sole on the right, and the Atlantic halibut (*Hippoglossus hippoglossus*) genome (GCF_009819705.1) on the left**. Crossing lines indicate chromosomal rearrangements.

### Genome assembly and polishing

We assembled the genome of the lemon sole using Hifiasm v0.19.5-r587 [10] with standard parameters. The resulting genome was analyzed using Inspector v1.0.1 [11], reporting a total sequence length of 1.09 Gbp, 1,641 contigs and an N50 of 12.18 Mbp. The mapping rate was 90.15 % and the QV was 38.78. The assembly was afterwards polished three times with Inspector using default parameters for HiFi reads.

### Assembly QC and scaffolding

To achieve chromosome-length scaffolds, we used the long-read based assembly and the generated HiC data as input for YaHS v1.1 [12] and Chromap v0.2.5 [13]. We mapped the HiC reads to the reference genome in HiC mode in Chromap, sorted the resulting SAM file, and converted it into the BAM format. Subsequently, this BAM file was used as input for YaHS with default settings. The assembly was initially scaffolded into 23 chromosome-length scaffolds and a total length of 1.09 Gbp. After manual curation with JuiceBox v2.20.00 [14], we obtained 24 chromosome-length scaffolds (Figure 3C). We filtered all scaffolds not containing any BUSCO sequences, as the size and fragmentation suggested assembly artifacts. The Contact map was created with HapHiC [15] using the initial HiC BAM file from Chromap and the final assembly graph (AGP).

Afterwards, the genome was gap filled with TGS Gapcloser v1.2.1 [16] set to PacBio reads and the “-x map-hifi” flag. The finalized assembly contained 38 scaffolds (24 chromosome-level) with a total length of 628 Mbp, a scaffold N50 of 27.21 Mbp, and an L50 of 11 (Figure 1B).

We evaluated both the raw and the final assembly with Merqury v1.3 in combination with Meryl v1.4.1 [17], as well as Inspector. We calculated assembly contiguity statistics of the final genome using QUAST v5.0.2 [18] and performed a gene set completeness analysis using compleasm v0.2.6 [19] with the provided Actinopterygii orthologous genes database (actinopterygii_odb10). The final genome was assembled to a total length of 628 Mbp with a scaffold N50 of 27.21 Mbp and a scaffold L50 of 11. The gene set completeness analysis identified 99.89 % complete BUSCO genes (98.62 % complete and single-copy) and only 0.08 % missing BUSCOs, which suggests that the assembly contains the majority of the genes expected to be found in Actinopterygii species (Fig. 1B). The Merqury analysis of the raw assembly showed a k-mer assembly completeness of 90 % and a quality value of 60.1. The final assembly resulted in an assembly completeness of 89.4 % and a quality value of 56. The resulting decrease is explained by scaffolding and removal of the assembly artifacts. The Inspector analysis of the final genome showed a mapping rate of 97.79 % and a depth of 97.04x. The Inspector QV was calculated to be 39.1 (Table S1).

Genome synteny of the final assembly was visualized using Jupiterplot v1.1 [19] with the Atlantic halibut assembly as reference (*Hippoglossus hippoglossus;* GCF_009819705.1) [20] (Fig. 1D). In both genomes, the Jupiterplot analysis was limited to the 24 chromosome-level scaffolds.

### Genome annotation

#### Repeat annotation

To annotate repetitive regions in the lemon sole genome, we used Earl Grey v6.0.1 [21], which incorporates RepeatModeler v2.0.16 [22] and RepeatMasker v4.1.5 [23]. EarlGrey was run with species set to Pleuronectidae using RepBase RepeatMasker Edition [24] and DFam v3.7 [25]. In total, 27.61 % of the genome consists of repeats (Table 1).

**Table 1.** Repeat content of the lemon sole genome assembly. Repeat content summary with TE class statistics found in the genome.

| TE Classification | Coverage (bp) | Copy Number | % Genome Coverage | Genome Size | TE Family Count |
| --- | --- | --- | --- | --- | --- |
| DNA | 59758564 | 256592 | 9.509771377 | 628391174 | 822 |
| Rolling Circle | 5929389 | 34674 | 0.9435824762 | 628391174 | 38 |
| Penelope | 745793 | 1597 | 0.1186829209 | 628391174 | 11 |
| LINE | 34470206 | 108384 | 5.48546947 | 628391174 | 331 |
| SINE | 3162509 | 21212 | 0.5032707541 | 628391174 | 25 |
| LTR | 14962171 | 44206 | 2.381028191 | 628391174 | 145 |
| Other (Simple Repeat, Microsatellite, RNA) | 21705113 | 337912 | 3.45407668 | 628391174 | 10044 |
| Unclassified | 32814865 | 214530 | 5.222044223 | 628391174 | 629 |
| Non-Repeat | 454842564 | NA | 72.38207391 | 628391174 | NA |

After the repeat analysis we hardmasked only complex, interspersed repeats (option: **-nolow**) from the resulting repeat library with RepeatMasker.

#### Gene annotation

Gene annotation was performed on the hardmasked assembly using GeMoMa v1.9 [26] with annotated high-quality genomes of other available flatfish genomes as reference (Table S2). The annotation resulted in 22,713 Genes, and 42,982 transcripts. Gene set completeness analysis based on the actinopterygii_odb10 database of the final genome annotation identified 98.5 % complete BUSCOs (62.99 % complete and single-copy, 33.79 % duplicated), 1.07 % fragmented BUSCOs, and 2.14 % missing BUSCOs, suggesting a high annotation completeness.

## Conclusion

Here, we report a chromosome-level genome assembly of the lemon sole (*Microstomus kitt*). The size of the scaffolded and annotated genome is in the range of other Pleuronectidae species and the number of scaffolds is also consistent with the haploid number of chromosome-length scaffolds from assemblies of other Pleuronectidae species [20,27]. With its high completeness and continuity, this genome provides important reference data for future population genetics analyses of this organism.

## DATA AVAILABILITY

All raw data generated in this study including PacBio HiFi long reads, HiC reads, and the chromosome-level assembly are accessible at GenBank under BioProject PRJNA1153561. Annotation, results files, and other data are available in the GigaDB repository.

## AUTHOR CONTRIBUTIONS

DP, MNe, MN, and AJ designed the study. DP, MN, MNe, AJ, JMOG, JJ, MMZ, FL, KW, HW, SCB, performed the laboratory work. DP, MN, MNe, AJ, JMOG, JJ, MMZ, FL, KW, HW, SCB, and MdJ conducted bioinformatic processing and analyses. All authors contributed to writing this manuscript.

### CONSENT FOR PUBLICATION

Not Applicable.

## ACKNOWLEDGEMENTS

We thank Alexander Ben Hamadou and Charlotte Gerheim from the TBG laboratory centre for laboratory support. The present study is a result of the Centre for Translational Biodiversity Genomics (LOEWE-TBG) and was supported through the programme ‘LOEWE – Landes-Offensive zur Entwicklung Wissenschaftlich-ökonomischer Exzellenz’ of Hesse’s Ministry of Higher Education, Research, and the Arts.

## SUPPLEMENTARY INFORMATION

**Supplementary Table 1.** Inspector quality metrics from the final genome assembly.

| Metric | Value |
| --- | --- |
| Number of contigs | 38 |
| Number of contigs > 10000 bp | 38 |
| Number of contigs >1000000 bp | 27 |
| Total length | 628391174 |
| Total length of contigs > 10000 bp | 628391174 |
| Total length of contigs >1000000bp | 622331302 |
| Longest contig | 34310146 |
| Second longest contig length | 32089857 |
| N50 | 27211925 |
| N50 of contigs >1Mbp | 27211925 |
| Mapping rate (%) | 97.79 |
| Split-read rate (%) | 14.09 |
| Depth | 97.0413 |
| Mapping rate in large contigs (%) | 97.22 |
| Split-read rate in large contigs (%) | 14.14 |
| Depth in large conigs | 97.417 |
| Total | 45 |
| Expansion | 22 |
| Collapse | 9 |
| Haplotype switch | 10 |
| Inversion | 4 |
| Errors per Mbp | 12.42697276 |
| Total small-scale assembly error | 7809 |
| Base substitution | 6568 |
| Small-scale expansion | 749 |
| Small-scale collapse | 492 |
| QV | 39.13441022 |

**Supplementary Table 2.** Available flat fish assemblies, used as reference for genome annotation with GeMoMa.

| Species | Accession Number |
| --- | --- |
| Common dab<br>(*Limanda limanda*) | GCF_963576545.1 |
| European flounder<br>(*Platichthys flesus*) | GCF_949316205.1 |
| European plaice<br>(*Pleuronectes platessa*) | GCF_947347685.1 |
| Turbot<br>(*Scophthalmus maximus*) | GCF_022379125.1 |
| Common sole<br>(*Solea solea*) | GCF_958295425.1 |

